# EARLY NEUTROPHIL ACTIVATION AND NETs RELEASE IN THE PRISTANE-INDUCED LUPUS MICE MODEL

**DOI:** 10.1101/2024.06.05.597651

**Authors:** Solange Carrasco, Bernadete L. Liphaus, Tatiana Vasconcelos Peixoto, Thais Martins Lima, Sueli Kunimi Kubo Ariga, Zelita Aparecida Jesus Queiroz, Thays de Matos Lobo, Sergio Catanozi, Letícia Gomes Rodrigues, Antônio Santos Filho, Walcy Rosolia Teodoro, Ana Paula Pereira Velosa, Débora Levy, Francisco Garcia Soriano, Cláudia Goldenstein-Schainberg

**Author notes:** Correspondence: Solange Carrasco, Laboratório de Imunologia Celular (LIM-17), Faculdade de Medicina, Universidade de São Paulo, São Paulo, Brazil.

## Abstract

**Background:** NETosis is recognized as an important source of autoantigens. Therefore, we hypothesized whether the pristane-induced lupus mice model shows early activation of neutrophils, the presence of low-density granulocytes (LDGs), and neutrophil extracellular traps (NETs) release, which could contribute to the development of a lupus phenotype.

**Methods:** Twelve female wild-type Balb/c mice were intraperitoneally injected with pristane (n=6; pristane group) or saline (n=6; control group). Five days after the injection, blood, peritoneal lavage, bone marrow, and spleen samples were collected for flow cytometry analyses of activated neutrophils (Ly6G+CD11b+), LDGs (CD15+CD14low), and NETs release (Sytox Green+).

**Results:** The pristane-induced mice group had a significantly increased number of blood activated neutrophils and LDGs as well as NETs released by these cells compared to the saline-injected control group and the basal values determined 12 days before the injection. The pristane group also had a significantly increased number of activated neutrophils, LDGs, and NETs released compared to the control group for the peritoneal lavage, bone marrow, and spleen.

**Conclusions:** We demonstrated early changes in the innate immune response such as an increased number of activated neutrophils and LDGs and mainly increased NETosis in the pristane-induced mice model which may be considered as the primary event triggering lupus development.

## Introduction

Pristane, a tetramethylpentadecane hydrocarbon oil, is well known for inducing in mice features of systemic lupus erythematosus (SLE) as autoantibodies, arthritis, peritonitis, and glomerulonephritis [1-4].

Born in bone marrow, neutrophils are innate immune first-line effector cells defense against invading pathogens [5-8]. Neutrophils are activated by a series of different PAMPs and DAMPs [9,10]. The CD11b and the Ly6G are neutrophil receptors responsible for: 1-cell migration, 2-making permanent connection with the endothelium, and 3-characterizing its phenotypic expressions [5,11]. CD11b is present on human and animal neutrophils, while Ly6G is merely observed in mice [5,11]. Notably, CD11b expression increases after neutrophils activation [5]. Both qualitative and quantitative alterations are observed in neutrophils from patients with autoimmune diseases as SLE [5,10,12-14]. These alterations include increased number of activated neutrophils [9], reduced phagocytosis capacity with lesser removal of apoptotic material [15-17], increased release of neutrophil extracellular traps (NETs), and increased number of low-density granulocytes (LDGs) [18-21].

LDGs are a subset of neutrophils, morphologically different, with a less segmented nuclei and a more lobular shape [18]. LDGs are characterized by INF type 1 secretion, proinflammatory activity, and enhanced spontaneous NETs release [18-22]. LDGs express neutrophil markers as CD15, and monocyte markers as CD14, suggesting these cells are immature neutrophils [21,23]. However, some authors believe LDGs arise from mature neutrophils that undergo degranulation after activation and, this way, are less dense [18,22]. Therefore, LDGs are considered a heterogeneous population of mature and immature neutrophils with a still discordant surface marker pattern [18,22]. During acute inflammation, activated neutrophils and LDGs were observed in primary and secondary lymphoid organs, and peripheral blood of SLE patients [20,24]. Increased amounts of LDGs in SLE patients have been related to disease activity and cutaneous involvement [18-21,23].

Our research group recently observed that the pristane-induced lupus mice model show increased number of activated T lymphocytes and reduced number of regulatory T cells (Treg) [25]. However, the innate immune response in this lupus model needs to be better understood.

Therefore, we hypothesize whether the pristane-induced lupus mice model show early activation of neutrophils, the presence of LDGs, and NETs release which could contribute to the development of a lupus phenotype.

## Methods

### Mice

Twelve female wild-type Balb/c mice, 8 to 10 weeks old, were provided by the Multidisciplinary Center for Biological Research – CEMIB/UNICAMP (Campinas, Brazil), and they were maintained in the Rheumatology Division, University of São Paulo School of Medicine - Brazil. All animal protocols were approved by the Institutional Animal Care and Research Advisory Committee (CAPPesq HC-USP Protocol # 1016-18), and conducted according to the U.S. National Institutes of Health (NIH) Guide for the Care and Use of Laboratory Animals. Mice were kept in an acclimatized facility with an automatic dark side cycle (http://www.biot.fm.usp.br/) and food and water were available *ad libitum*. After the acclimatization period, at 10 to 12 weeks of age, and 12 days before the intraperitoneal injection, the time required for recovery of the leukocyte pool, a peripheral blood sample was collected from all animals. This blood sample was used as the normal or basal number of activated neutrophils, LDGs, and NETs released by both the former cells. Mice used in the experiments weighed 20 to 25g, and they received a single intraperitoneal injection of 0.5 ml of pristane (TMPD-tetramethylpentadecane Sigma Chemical Co., St. Louis, MO) (n=6; pristane group), as we previously used for SLE induction [25], or an equal volume of 0.9% saline (n=6; control group). Five days after pristane or saline was intraperitoneally injected, that is, the shortest period ever analyzed according to our literature review [1-4], and before the disappearance of neutrophils after the inflammation induction, samples of blood, peritoneal lavage, bone marrow, and spleen were collected from both mice groups for flow cytometry analysis.

### Blood samples

Immediately after collecting, 200 µL of peripheral blood from the caudal vein 12 days before the intraperitoneal injection, and 500 µL of blood from the inferior vena cava five days after pristane or saline injection, red blood cells were lysed by incubating samples with FACSTM Lysing Solution (Becton Dickinson New Jersey NJ USA). After that, samples were centrifuged (800 g for 10 min at 4ºC), the cells pellet washed, and it was suspended in RPMI containing 5% fetal bovine serum (FBS). The hemolysis method was chosen as it has simple steps and preserves all types of white blood cells (granulocytes, monocytes, and lymphocytes). Cell suspensions were promptly analyzed by flow cytometry.

### Peritoneal lavage sample

Five days after pristane or saline injection, mice were euthanized with anaesthetic overdose, 2 mL of RPMI were intraperitoneally injected, and the cell suspension recovered. The peritoneal lavage was centrifuged (800 g for 10 min at 4ºC), the cells pellet washed, and it was suspended in RPMI containing 5% FBS. The cell suspension, containing all types of white cells was counted in a Neubauer chamber using trypan blue (1:1) to ensure a differential count of at least 1×10^6^ polymorphonuclear and mononuclear cells.

### Bone marrow sample

Immediately after mice were euthanized, femurs and tibias were obtained and the bone marrow was washed with Hank’s balanced salt solution with EDTA. The recovered suspension was centrifuged (800 g for 10 min at 4ºC), the cells pellet washed, and it was suspended in RPMI containing 5% FBS. The bone marrow cell suspension was promptly analyzed by flow cytometry.

### Spleen sample

After mice euthanasia, the spleen was removed, chopped, and crushed in a plate with RPMI medium. The cell suspension was transferred to a Falcon tube, which stood for approximately 2 min for precipitation of larger tissue samples. Red blood cells were lysed by incubating samples with lysing solution. Samples were then centrifuged (800 g for 10 min at 4ºC), and the cell pellet was washed twice and suspended in RPMI containing 5% FBS. The cell suspension was counted in the Neubauer chamber to obtain at least 1×10^6^ polymorphonuclear and mononuclear cells.

### Monoclonal antibodies

The following anti-mouse monoclonal antibodies (BD Biosciences New Jersey NJ USA) were used: Ly6G PE-Cy7 (clone:1A8), Ly6G PE (clone:1A8), CD14 PE (clone:mC5-3), CD11b APC-Cy7, (clone:M1/70), and Anti-SSEA-1 APC-Cy5.5 CD15 (clone:MC480). Sytox Green from Invitrogen was also used. Isotypic controls were used according to the company’s protocols.

### NETs release assays

NETs released by activated neutrophils (Ly6G+CD11b+) and LDGs (CD15+CD14low) were quantitatively measured by flow cytometry [26]. NETs release was detected by staining the cells with the fluorescent dye for DNA Sytox Green, which does not permeate the cell membrane, and bind to pendant-DNA [26]. The number of activated neutrophils and LDGs in NETosis was determined by flow cytometry using triple staining (Ly6G+CD11b+Sytox+ and CD15+CD14lowSytox+, respectively). Additionally, NETs released by neutrophils were qualitatively demonstrated by the high-content screening technique. Sytox Green is reliable for detection of NETs by both flow cytometry and fluorescent staining [26].

### Flow cytometry assays

Cells from blood, peritoneal lavage, bone marrow, and spleen were stained with the Ly6G, CD11b, CD15, and CD14 monoclonal antibodies in the dark for 30 min at 4 to 8ºC. After this period, phosphate-buffered saline (PBS)+SFB was added, and cells were centrifuged (800g for 5 min at room temperature). The supernatant was discarded, cells were suspended with Sytox Green+PBS+SFB, and incubated in a CO_2_ ambient for 30 min at 37ºC. The multiparameter flow cytometry assays were carried out using Guava EasyCyteTM HT (Millipore Minneapolis, NM, USA), and the analyses were conducted using InCyte software (Millipore). Based on forward and side scatter, an R1 region containing the neutrophil and the monocyte populations was determined. Within the R1 region, activated neutrophils were detected on a double-positive Ly6G and CD11b plot (R2 region). Within the R2 region, Sytox Green-positive activated neutrophils were quantified for NETs release. In addition, within the same R1 region, LDGs were detected on a double-positive CD15 and CD14low plot (R3 region), since these are the surface markers most related to autoimmune diseases in literature [12]. Within the R3 region, Sytox Green-positive LDGs were quantified for NETs release.

### High-content screening assay

Cell suspensions of blood, peritoneal lavage, bone marrow, and spleen were diluted with 3 mL of PBS containing 0.5% bovine albumin. Each diluted suspension was submitted to the Ficoll-Hypaque 1.077 and 1.119 gradient technique. After collecting the interface between gradients, 1×10^4^ neutrophils were distributed on a plate, and they were stained with Ly6G monoclonal antibody which colored them in red. Cell suspension was then stained with Sytox Green which highlights NETs in green. Finally, the plate was stained with Hoechst dye, which colored nuclei in blue.

### Statistical analysis

Data are expressed as mean ± SD as well as median, minimum, and maximum. Groups were compared using generalized estimation equations with normal or Poisson distribution and identity link function assuming an interchangeable correlation between locations. Results were followed by Bonferroni multiple comparisons to identify differences between groups. P values ≤ 0.05 were considered statistically significant.

## Results

### The pristane-induced mice model shows an increased number of activated neutrophils

The pristane-induced mice group had a significantly increased number of activated neutrophils (Ly6G+CD11b+) of blood compared to both the saline-injected mice group (control) (4,120.2 ± 2,648.5 vs 1,149 ± 183.8, *p* < 0.001) as well as the basal value (the sample collected 12 days before the pristane injection) of activated neutrophils (4,120.2 ± 2,648.5 vs 660.6 ± 478.3, *p* < 0.001). An increased number of activated neutrophils was also observed in the pristane group compared to the saline group for the samples of peritoneal lavage (4,953 ± 797 vs 569.7 ± 538.4, p < 0.001), bone marrow (6,222.8 ± 515.3 vs 302.3 ± 48.3, p <0.001), and spleen (2,119.7 ± 638.6 vs 1,739 ± 102.5, *p* < 0.001), as shown in Figure 1.

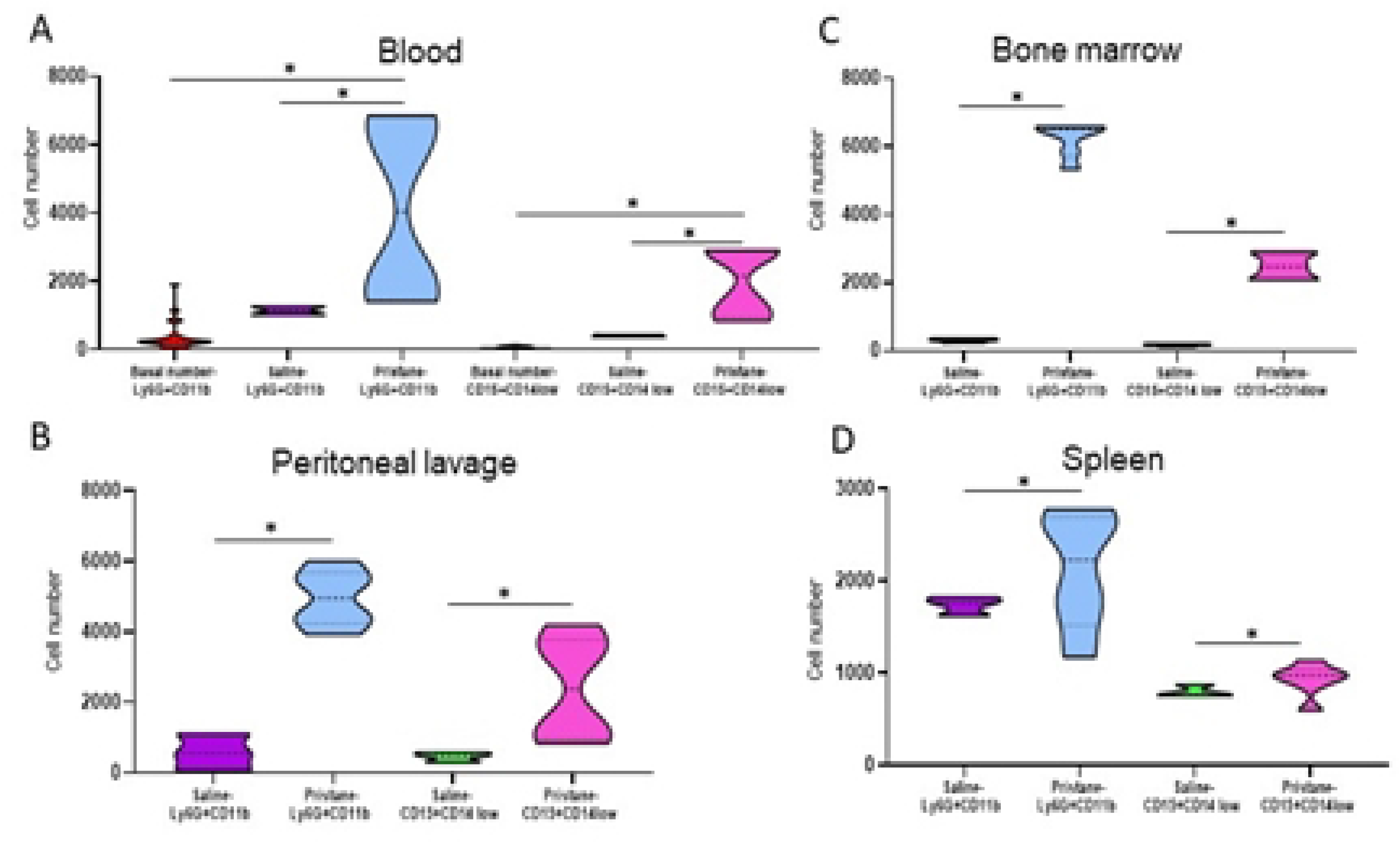

### The pristane-induced mice model shows an increased number of LDGs

The pristane group had a significantly increased number of LDGs of blood compared to both the saline group (1,998.3 ± 1,012.4 vs 418 ± 36.8, *p* < 0.001) and the basal value of LDGs (1,998.3 ± 1,012.4 vs 38.8 ± 37.7, *p* < 0.001). An increased number of LDGs was also observed in the pristane group compared to the saline group for the samples of peritoneal lavage (2,813 ± 1,032.8 vs 448.3 ± 141.5, *p* < 0.001), bone marrow (2,483.5 ± 372.2 vs 168.7 ± 25.9, *p* < 0.001), and spleen (929.2 ± 182.7 vs 802 ± 65.2, *p* < 0.001), as shown in Figure 1.

### The pristane-induced mice model shows an increased NETs release by activated neutrophils

The pristane group had a significantly increased NETs release by activated neutrophils of blood compared to the saline group (3,371.5 ± 2,162.9 vs 393 ± 49.5, *p* < 0.001), and the basal value of NETs release (3,371.5 ± 2,162.9 vs 383.2 ± 423.7, *p* < 0.001). An increased NETs release by activated neutrophils was also observed in the pristane group compared to the saline group for the samples of peritoneal lavage (2,035.5 ± 636.1 vs 308.7 ± 285.9, *p* < 0.001), bone marrow (1,735 ± 639.7 vs 154 ± 29.1, *p* < 0.001), and spleen (1,179 ± 299 vs 432.7 ± 113.8, *p* < 0.001), as shown in Figure 2. Representative images of the high-content screening immunostaining of NETs release by activated neutrophils in all the evaluated sites Figure 3.

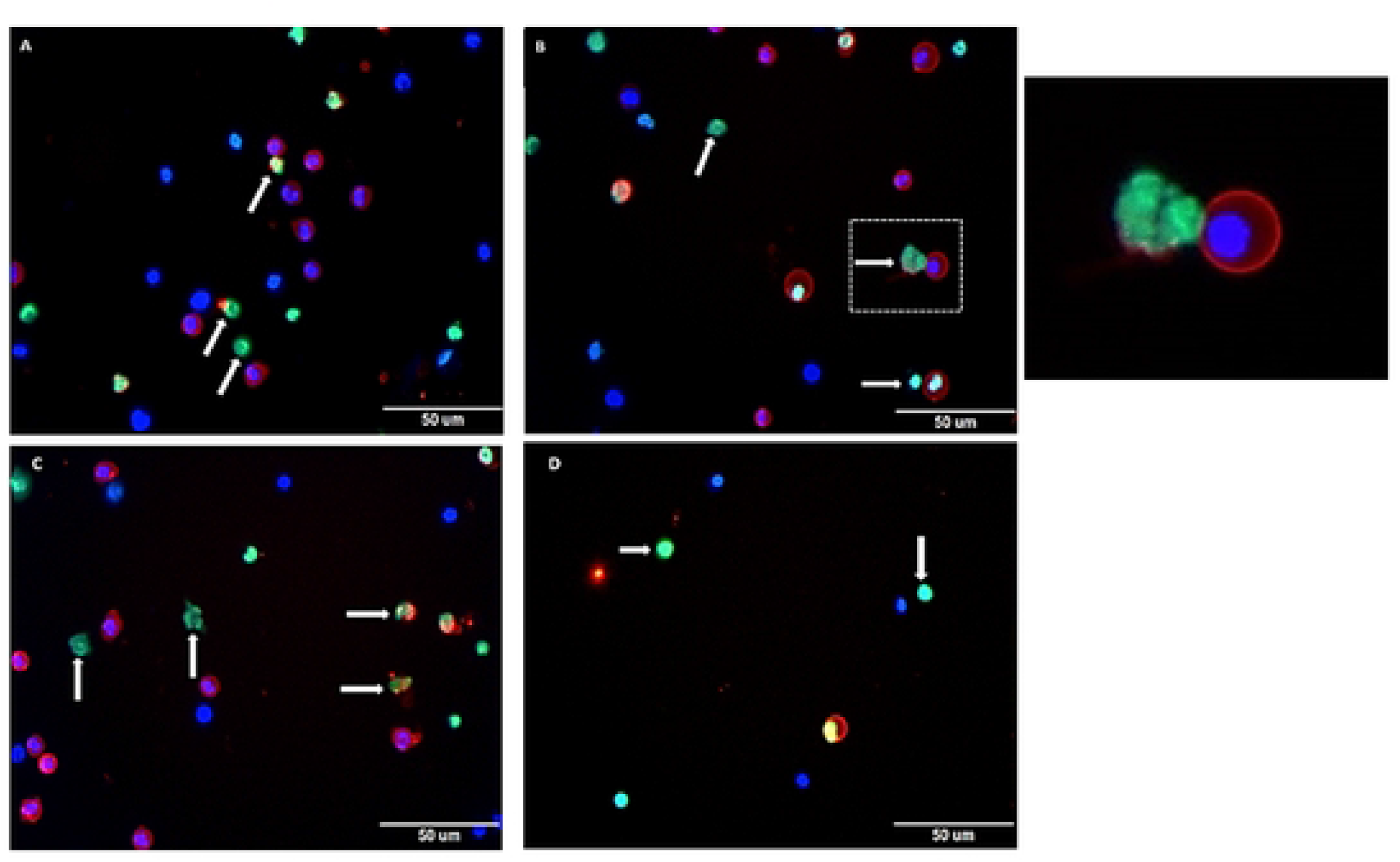

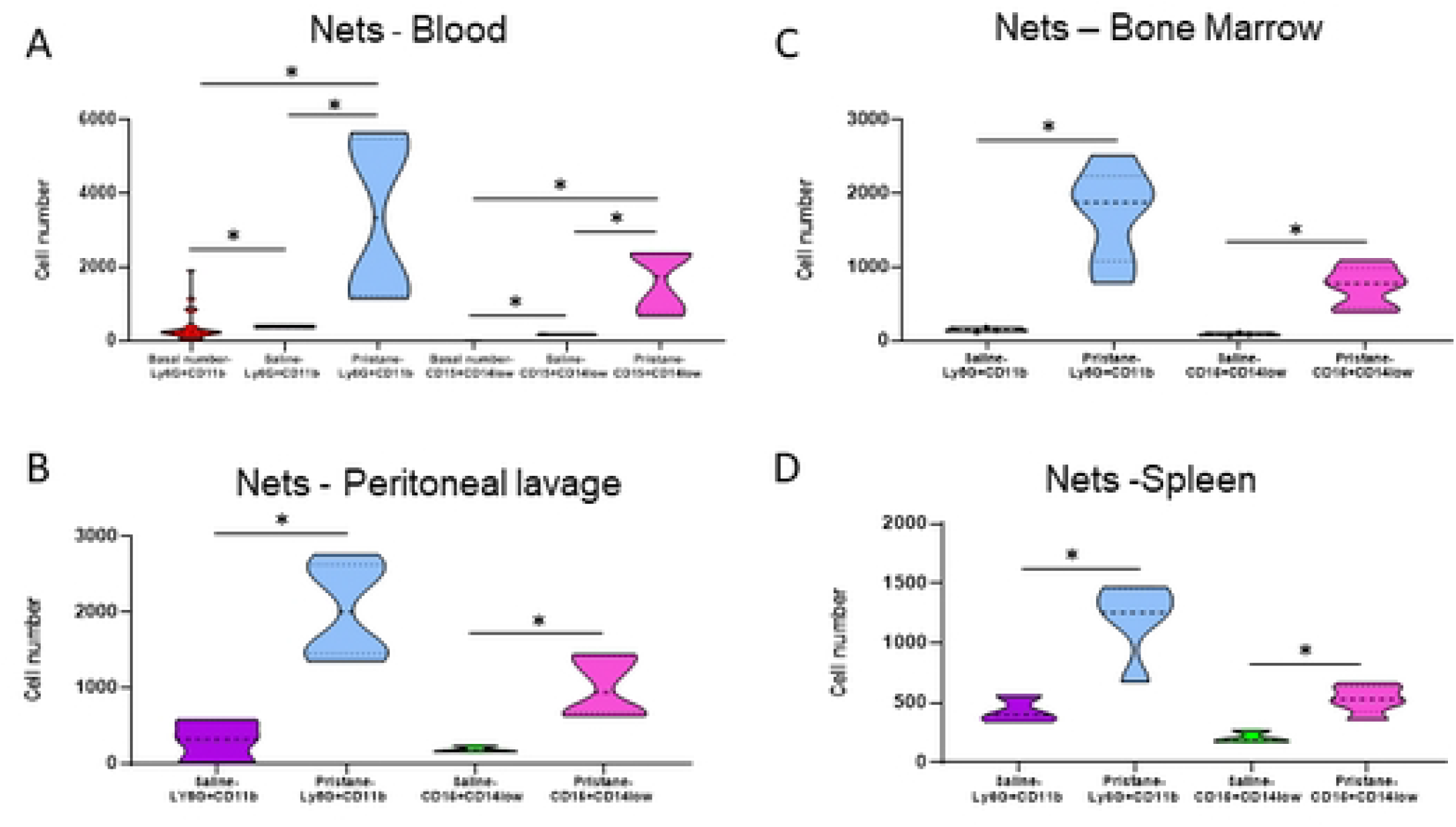

### The pristane-induced mice model shows an increased NETs release by LDGs

The pristane group had significantly increased NETs release by LDGs of blood compared to the saline group (1,623.3 ± 824.3 vs 178 ± 29.7, *p* < 0.001), and the basal value of NETs released by LDGs (1,623.3 ± 824.3 vs 5.7 ± 6.8, *p* < 0.001). An increased NETs release by LDGs was also observed in the pristane group compared to the saline group for the samples of peritoneal lavage (1,011 ± 387.5 vs 186.7 ± 42.2, *p* < 0.001), bone marrow (744.5 ± 275 vs 93 ± 20.1, *p* < 0.001), and spleen (534 ± 106.4 vs 209.7 ± 52.4, *p* < 0.001), as shown in Figure 2. Results are also summarized in the supplementary table.

## Discussion

This study shows that the pristane-induced mice model present as early as five days increased number of activated neutrophils and LDGs, accompanied by increased NETs release by these cells in all the sites evaluated namely, blood, peritoneum, bone marrow and spleen, corroborating the role of the innate immune system in the development of lupus.

In human lupus, it has been shown that large amounts of intracellular debris exposed during the acute inflammatory process by neutrophils act as the major source of autoantigens [10,12,13,18,27-29]. In addition, LDGs are present in increased amounts and related to disease activity [18-21,23] and cutaneous involvement [18,20,21]. Recently, the inhibitory action of gasdermin D on LDGs from patients and the pristane-induced mice model was demonstrated, since disease severity was reduced with this treatment [30]. In this regard, our study shows that not only an increased number of LDGs occur in lupus, but also increased NETs released by these cells. This study also shows that the former alterations occur not only in blood, but also in the peritoneum, and the lymphoid organs such as bone marrow and spleen.

NETs are rich in DNA and histones and participate in the presentation of autoantigens, inducing increased production of pro-inflammatory cytokines during the immune response [10,14,20,28,31,32]. In patients with SLE, increased amounts of NETs in peripheral blood have also been described, similar to our findings in mice [13,14,20,32]. Furthermore, impaired clearance of NETs results in increased anti-NET and anti-dsDNA autoantibodies, and the development of kidney damage [10,14,28,32]. It is important to emphasize that in SLE patients, neutrophils and LDGs have a great propensity to release NETs and undergo an accelerated process of apoptosis *in vitro*, as observed herein [18,30,33]. Thus, our results indicate that the primary and initial event for the autoantibodies production may be the early NETs release, as observed five days after the pristane injection. Therefore, the search for factors and procedures that block NETs release may be a way to prevent the development of lupus.

Lupus pictures are established in mice models in a median of 90 days after pristane injection and show massive autoreactive infiltration and cell death overwhelming the clearance process [25]. We observed an increased amount of activated T lymphocytes (CD4+CD69+) in peripheral blood and spleen [25]. However, it was remarkable that no T lymphocytes were found in the peritoneum, possibly due to cell migration from the application area to blood and other organs [25]. On the other hand, in this study, we observed an increased number of activated neutrophils and LDGs in all analyzed sites, blood, peritoneum, bone marrow, and spleen. These findings highlight the role of the innate immune system, neutrophils, and LDGs, with subsequent NETs release, as possible early alterations involved in the etiopathogenesis of lupus [17-21,23,24]. Few studies have analyzed NETs and LDGs in different animal models (BALB/cJ, galectin-3 deficient mice, wild-type C57BL/6, PAD4, LysMCre-PAD4, pristane-induced, Gsdmd^-/-^ mice, WT, and rats) and periods (seven, fourteen days, three and seven months) [3,4,30,31,34,35] but no research at an early period and with so many different anatomic sites as in this study. It is interesting to note that five days after pristane injection the immune response is reasoned on activated neutrophils and not on the acquired immune system producing antibodies. Lymphocytes will be induced to produce autoantibodies late in the process of lupus disease. NETs release is an extreme action of neutrophils against invaders due to the huge stimulus loaded by PAMPs [6]. This data appears to accompany the increase in the number of LDGs, which may represent activated neutrophils that have already released their cytoplasmic granules, and as already activated cells, due to possible pre-exposure to PAMPs, their next action can only be NETs release [6,7,23]. Again, this data shows that the beginning of the lupus induction process passes through the innate immune system and only later reaches the acquired immune system.

Another relevant aspect identified in this study is the fact that the immune response is systemic, that is, after a local injection of pristane into the peritoneum, activated neutrophils, LDGs, and NETs release were identified not only locally, but also in blood, bone marrow, and spleen. This systemic behavior corresponds to infectious processes that evolve into sepsis, a systemic condition [6,7]. It is relevant to note that lupus patients have a systemic condition with the impairment of several organs due to autoimmune activity [22].

To the best of our knowledge, this is the first study to analyze, in mice model, the pristane induction effects at the bone marrow, the primary site of immune cell production. Similar to what happens in the peritoneum, an acute increase in the number of activated neutrophils and LDGs, as well as an enhanced NETs release by these cells were demonstrated in the bone marrow.

The strong points of present study were: 1-the use of fresh cells labeled right after their isolation which avoids any spontaneous activation or NETs release in contrast to studies that used frozen cells; 2-neutrophils and LDGs changes were analyzed right after the breakdown of the immune tolerance (five days after pristane injection); 3-NETs release was assessed in both neutrophils and LDGs; 4-neutrophils and LDGs changes were evaluated at different anatomic sites including the primary injury site (peritoneum), the circulation (blood) and the lymphoid organs (bone marrow and spleen).

This study limitations were: 1-the small number of samples analyzed; 2-the lack of evaluating the behavior of neutrophils and LDGs over different periods; and 3-not distinguishing the LDGs subsets.

In conclusion, we demonstrated early changes in the innate immune response such as an increased number of activated neutrophils and LDGs and mainly increased NETosis in the pristane-induced mice model which may be considered as the primary event triggering lupus development.

## Acknowledgments

The authors acknowledge Fundação de Amparo à Pesquisa do Estado de São Paulo (FAPESP) for the financial supported by grants 2017/02335-0 and 2013/19292–1.

---

**Supplementary Table.** Number of activated neutrophils, low-density granulocytes, and NETs release by activated neutrophils and low-density granulocytes by each mice group and site.

| Site / Group | Basal value (n= 12) | Saline (n=6) | Pristane (n=6) |
| --- | --- | --- | --- |
| **Activated neutrophils** |  |  |  |
| **Peripheral blood** | 660.6 ± 478.3<br>534 (86 – 1,981) | **1,149 ± 183.8\***<br>1,149 (1,019 – 1,279) | **4,120.2 ± 2,648.5\*§**<br>4,043 (1,411 – 6,862) |
| **Peritoneal lavage** |  | 569.7 ± 538.4<br>547 (43 – 1,119) | **4,953 ± 797§**<br>4,947 (3,926 – 6,005) |
| **Bone marrow** |  | 302.3 ± 48.3<br>315 (249 - 343) | **6,222.8 ± 515.3§**<br>6,505 (5,332 – 6,590) |
| **Spleen** |  | 1,739 ± 102.5<br>1,762 (1,627 – 1,828) | **2,119.7 ± 638.6§**<br>2,230.5 (1,173 – 2,778) |
| **Low-density granulocytes** |  |  |  |
| **Peripheral blood** | 38.8 ± 37.7<br>31.5 (0 - 123) | **418 ± 36.8\***<br>444 (392 - 613) | **1,998.3 ± 1,012.4\*§**<br>2,162.5 (839 - 2924) |
| **Peritoneal lavage** |  | 448.3 ± 141.5<br>476 (295 - 574) | **2,813 ± 1,032.8§**<br>2,386.5 (839 - 4197) |
| **Bone marrow** |  | 168.7 ± 25.9<br>180 (139 - 187) | **2,483.5 ± 372.2§**<br>2,472.5 (2,079 – 2,922) |
| **Spleen** |  | 802 ± 65.2<br>770 (759 - 877) | **929.2 ± 182.7§**<br>976.5 (597 - 1140) |
| **NETs release by activated neutrophils** |  |  |  |
| **Peripheral blood** | **383.2 ± 423.7**<br>244 (56 – 1,925) | **393 ± 49.5**<br>393 (358 - 428) | **3,371.5 ± 2,162.9\*§**<br>3,350 (1,145 – 5,660) |
| **Peritoneal lavage** |  | 308.7 ± 285.9<br>327 (14 - 585) | **2,035.5 ± 636.1§**<br>2,007 (1,352 – 2,767) |
| **Bone marrow** |  | 154 ± 29.1<br>152 (126 - 184) | **1,735 ± 639.7§**<br>1,884.5 (789 - 2,523) |
| **Spleen** |  | 432.7 ± 113.8<br>397 (341 - 560) | **1,179 ± 299§**<br>1,245.5 (675 - 1471) |
| **NETs release by low-density granulocytes** |  |  |  |
| **Peripheral blood** | **5.7 ± 6.8**<br>2.5 (0 - 26) | **178 ± 29.7**<br>178 (157 - 199) | **1,623.3 ± 824.3\*§**<br>1,760 (676 - 2,388) |
| **Peritoneal lavage** |  | 186.7 ± 42.2<br>168 (157 - 235) | **1,011 ± 387.5§**<br>942 (637 – 1,448) |
| **Bone marrow** |  | 93 ± 20.1<br>91 (74 - 114) | **744.5 ± 275§**<br>778 (390 – 1,100) |
| **Spleen** |  | 209.7 ± 52.4<br>183 (176 - 270) | **534 ± 106.4§**<br>538.5 (359 - 666) |

Results are shown by mean, standard deviation, median, minimum, and maximum. \* $p < 0.001$ compared to the basal value ; § $p < 0.001$ compared to the saline group.

